# Microtubules restrict F-actin polymerization to the immune synapse via GEF-H1 to maintain polarity in lymphocytes

**DOI:** 10.1101/2022.01.03.473915

**Authors:** Judith Pineau, Léa Pinon, Olivier Mesdjian, Jacques Fattaccioli, Ana-Maria Lennon Duménil, Paolo Pierobon

## Abstract

Immune synapse formation is a key step for lymphocyte activation. In B lymphocytes, the immune synapse controls the production of high-affinity antibodies, thereby defining the efficiency of humoral immune responses. While the key roles played by both the actin and microtubule cytoskeletons in the formation and function of the immune synapse have become increasingly clear, how the different events involved in synapse formation are coordinated in space and time by actin-microtubule interactions is not understood. Using a microfluidic pairing device, we studied with unprecedented resolution the dynamics of the various events leading to immune synapse formation and maintenance. Our results identify two groups of events, local and global dominated by actin and microtubules dynamics, respectively. They further highlight an unexpected role for microtubules and the GEF-H1-RhoA axis in restricting F-actin polymerization at the immune synapse to define the cell polarity axis, allowing the formation and maintenance of a unique competent immune synapse.

## Introduction

Cell polarization refers to the acquisition of a cell state characterized by the asymmetric distribution of cellular individual components, including molecules and organelles. It is critical for a multitude of cellular functions in distinct cell types and further controls cell-cell interactions. This particularly applies to lymphocytes, which rely on cell polarity to form a stereotyped structure called the immune synapse to communicate with antigen presenting cells (1–5). Immune synapses are not only instrumental for lymphocyte activation but also serve their effector functions, for example by facilitating the killing of infected or malignant cells by cytotoxic cells (6, 7). Understanding how immune synapses form has thus become a major challenge for cell biologists and immunologists for the last decade, yet many mechanistic questions remain unanswered. In particular, how immune synapses are maintained in time to serve sustained lymphocyte function and allow robust immune activation is poorly understood.

Immune synapse formation is accompanied by the reorganization of lymphocyte antigenic receptors and associated signaling molecules into a concentric structure that forms at the contact zone with antigen presenting cells (1, 3). The synapse allows the exchange of information (molecules and vesicles) between the two cells through tightly regulated exocytic and endocytic events (8). Signaling and trafficking at the immune synapse require deep rearrangements of both the lymphocyte actin and microtubule cytoskeletons (9). On one side, the actin cytoskeleton controls the organization of antigen receptor-containing micro-clusters for coordination between trafficking and signaling and further helps generating the mechanical forces that depend on the myosin II motor (10–13). On the other side, the microtubule cytoskeleton controls the recruitment of organelles at the immune synapse. This relies on centrosome re-orientation, leading to lymphocyte symmetry breaking and acquisition of a polarized cell state (14, 15). Although it is now clear that these events of actin and microtubule re-organization are instrumental for synapse formation, how they depend on each other and are coordinated to ensure proper and durable synapse function remains elusive.

There is growing evidence in the literature suggesting that the actin and microtubule cytoskeletons do not act independently of each other but indeed functionally and/or physically interact (16, 17). This is well-illustrated, for example, by the study of oocyte polarization in C. *Elegans* where polarization of intracellular organelles occurs in response to actomyosin contraction at one cell pole, which is in turn down-regulated upon centrosome recruitment (18). A crosstalk between actin and microtubules in lymphocytes was also recently highlighted by our work showing that clearance of branched actin at the centrosome is needed for its detachment from the nucleus and polarization to the synapse (19). However, whether the microtubule network in turn impacts on actin dynamics and immune synapse formation, function and maintenance has not been studied, in part because the tools to quantitatively monitor in time both local actin reorganization and microtubule re-orientation were not available so far. In this work we developed a microfluidic chamber to quantitatively analyze both the local and global events associated to immune synapse formation in time and space and establish their dependency on actin and microtubule cytoskeletons. Our results revealed that the microtubule network controls the polarized polymerization of F-actin at the interface between lymphocytes and antigen presenting cells, thereby allowing sustained formation of a unique and functional immune synapse.

## Results

### A microfluidic system for the systematic study of immune synapse formation

We aimed at understanding how local and global events of synapse formation were coordinated in space and time. As a model, we used B lymphocytes, which form immune synapses upon engagement of their surface B Cell Receptor (BCR) by cognate antigens presented at the surface of neighboring cells. *In vivo*, this cell-cell interaction takes place in lymphoid organs and is required for antigen extraction and activation of signaling pathways that later-on promote B lymphocyte differentiation into cells able to produce high-affinity antibodies (4, 20). Antigen extraction involves two modes: (1) an early mechanical mode that relies on actin-mediated forces at the synapse and (2) a late proteolytic mode that requires centrosome polarization to the synapse and subsequent lysosomes transport on microtubules and secretion of hydrolases into the extracellular milieu (14, 21, 22). It has been shown that mechanical antigen extraction occurs on deformable substrates while proteolytic extraction is used to extract antigen from stiff materials (22). The first pathway, when activated, inhibits the second one (22), suggesting a functional interaction between these actin- and microtubule-dependent events. However, the experimental systems used so far did not allow to reach a sufficient temporal resolution to quantitatively monitor the evolution of both cytoskeleton networks in 3D from the first instant of immune synapse formation.

To circumvent this problem, we built a microfluidics device based on an array of traps where antigen-coated oil droplets and B cells can be sequentially captured (Figure 1A, Supp Movie 1). Antigen-coated lipid droplets are a good 3D substrate to mimic antigen-loaded cells, as they allow antigen mobility at their surface (Figure 1B). Moreover, they are effectively stiff (see material and methods) and might thus also allow lysosome recruitment at the synapse and proteolytic antigen extraction. Chambers were imaged in 3D from the time of cell injection to capture the entire process of synapse formation. Droplets were functionalized either with a non-activating molecule (BSA, negative control) or an activating BCR ligand (F(ab’)_2_ anti-Mouse IgG, referred to as “antigen” from now on). Both ligands were grafted to the lipid droplet with fluorescent streptavidin to follow their accumulation dynamics at the droplet surface (Figure 1B-D, Supp Movie 2). Such an accumulation was exclusively observed upon engagement of the BCR with its ligand, BSA-coated droplets remaining homogeneously fluorescent (Figure 1E, F). This result indicates that this system can be used to reconstitute immune synapses, and in particular study the role of actin-microtubule interactions in the formation, function and maintenance of this structure.

**Fig. 1.**
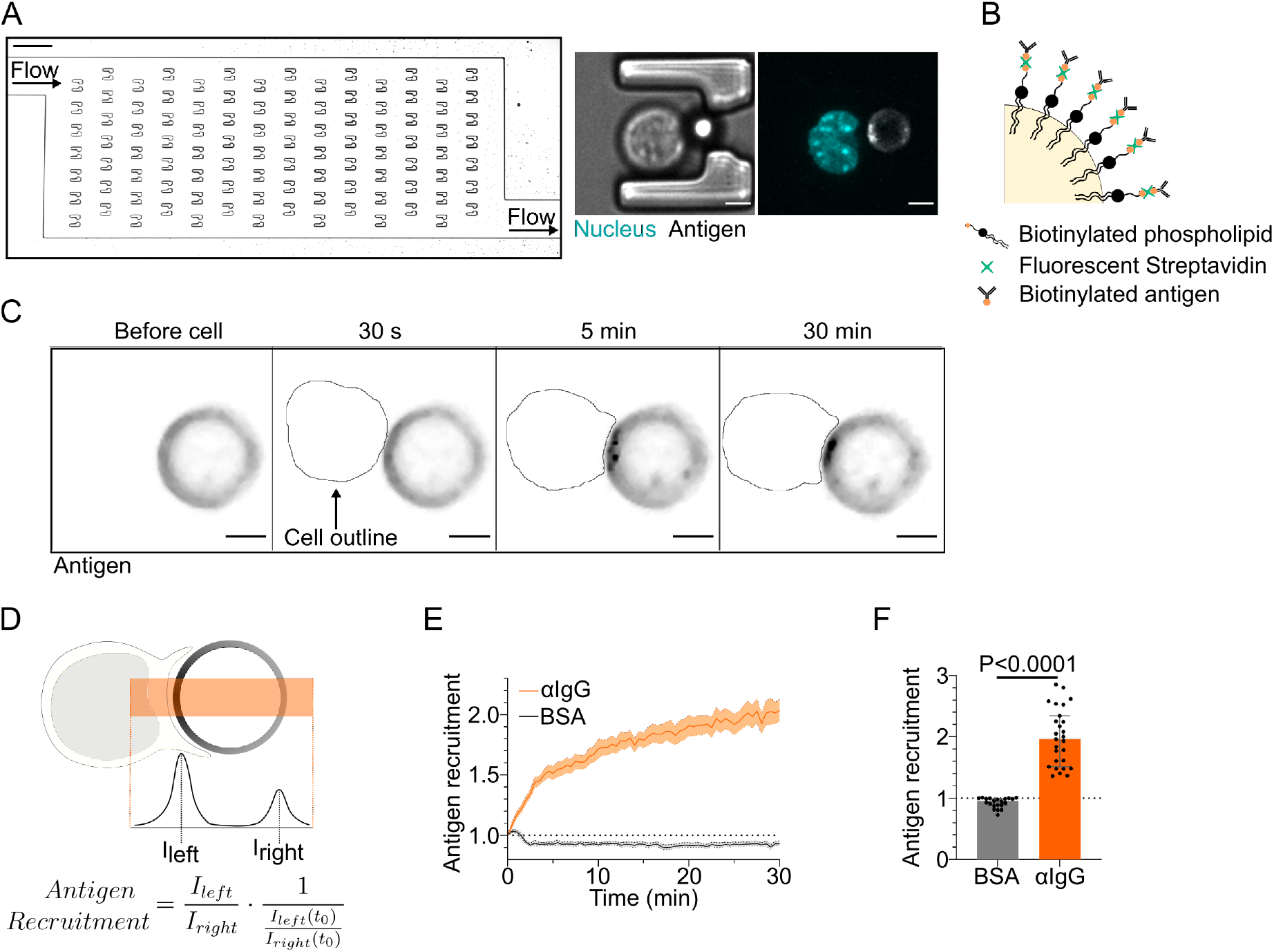
Microfluidic system to study dynamics of B lymphocyte polarization and immune synapse formation. (A) Transmission image of a chamber of the microfluidic chip containing the traps. Scale bar 100 μm. Inset: Cell-droplet doublet in a microfluidic trap. Bright field image and fluorescence image (Nucleus: cyan, Antigen: gray). Scale bar 5 μm. (B) Schematic representation of the surface of an oil droplet used for antigen presentation. (C) Time-lapse images of antigen recruitment on an F(*ab′*)_2_ *α*IgG-coated droplet (acting as an antigen). Scale bar 5 μm. (D) Schematic representation of the quantification of antigen recruitment at the immune synapse. (E) Quantification over time of recruitment on BSA-coated (negative control) or *α*IgG-coated droplets at the immune synapse (Mean±SEM, BSA N=21, *α*IgG N=27, 2 independent experiments). (F) Plateau of Antigen recruitment (average value 25-30 min) on BSA- or *α*IgG-coated droplets (Median±IQR, BSA N=21, *α*IgG N=27, 2 independent experiments, Mann-Whitney test).

### Defining characteristic timescales of immune synapse formation

Our microfluidic system was used at first to visualize and extract the typical timescales of the key events associated to synapse establishment: BCR signaling (production of DiAcylGlycerol (DAG) monitored by a GFP-C1*δ* reporter (23)), F-actin reorganization (labeled with F-tractin-tdTomato), centrosome (labeled with SirTubulin) and Golgi apparatus (labeled with Rab6-mCherry) polarization, lysosomes (labeled with Lysotracker) and nucleus (labeled with Hoechst) re-positioning. Characteristic timescales were extracted from volumetric images taken every 30 seconds (Supp Movie 3).

We found that the peak of DAG production occurred ~3.25 minutes upon contact between the lymphocyte and the antigen-coated droplet (Figure 2A, G). It was concomitant with actin polymerization, which peaked at the synapse at ~3 minutes (Figure 2B, G). Formation of the stereotypical actin pattern, with actin protrusions at the periphery and an actin-cleared area at the center, was then observed. Centrosome and Golgi tracking over time showed that they displayed similar behaviors, reaching the immune synapse area after 5 minutes for the centrosome (distance<2 μm) and 6.5 minutes for the Golgi apparatus (distance<4 μm) (Figure 2C, D, G). This was only observed in cells where the BCR was specifically engaged and is in good agreement with these two organelles being physically associated (24). Lysosomes, which are also known to associate with microtubules for intracellular transport, displayed a slightly different behavior: their distance to the immune synapse decreased down to ~3 μm in ~6 minutes, indicating their polarization, but then increased (Figure 2E, G), maybe due to the secretion of lysosomal vesicles which would lead to signal fainting at the immune synapse and a consequential apparent re-distribution all over the cell.

**Fig. 2.**
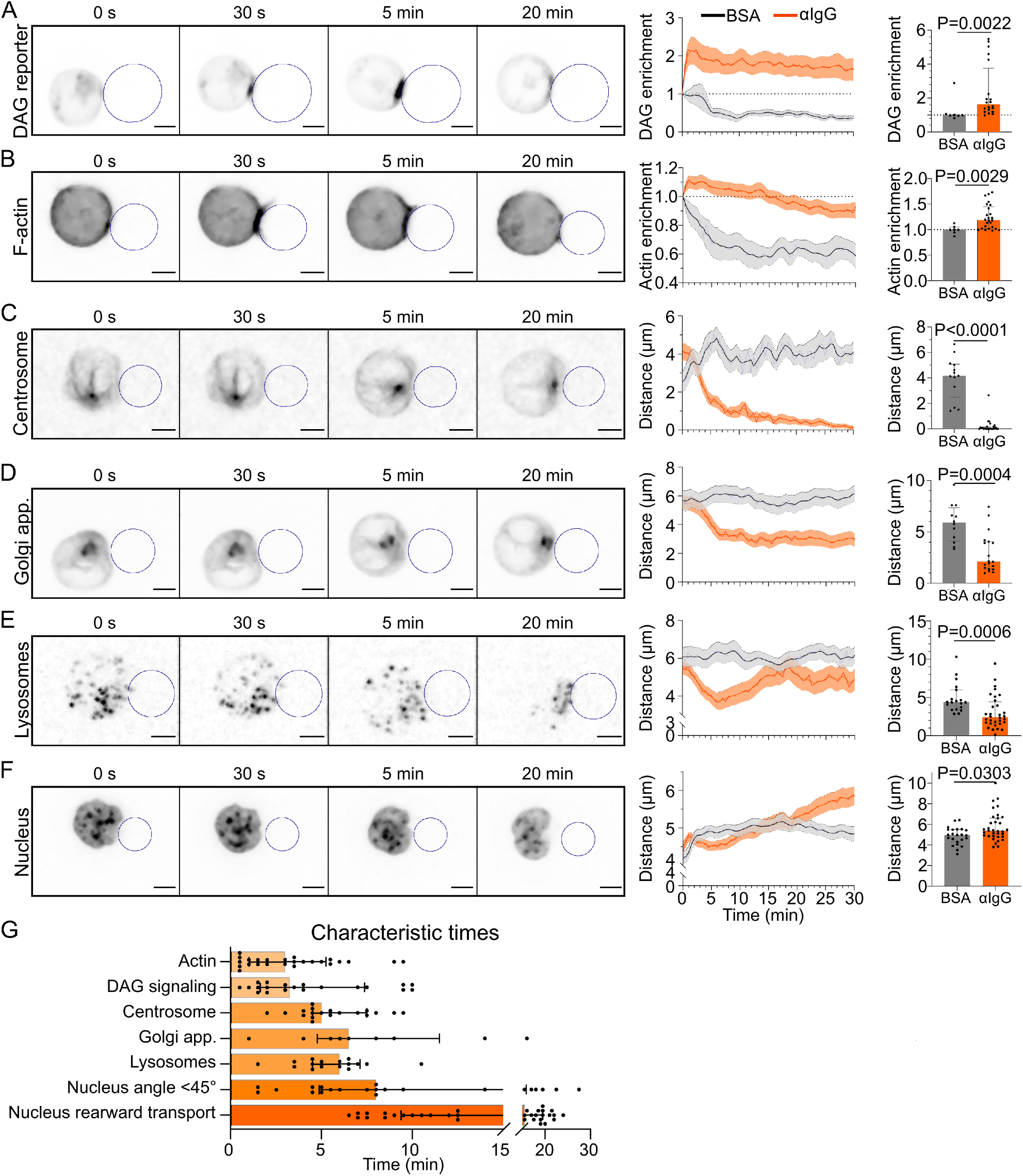
Timescales of B lymphocyte polarization. (A) Time lapse images of DAG reporter in a cell in contact with a droplet (outlined in blue). Enrichment in time of DAG reporter (Mean±SEM). Maximum enrichment (0-10 min), (Median±IQR, BSA N=7, *α*IgG N=20, 2 independent experiments, Mann-Whitney test). (B) Time lapse images of F-actin in a cell in contact with a droplet (outlined in blue). Enrichment in time of F-actin near the droplet for BSA- or *α*IgG-coated droplets (Mean±SEM). Maximum enrichment (0-10 min), (Median±IQR, BSA N=7, *α*IgG N=26, 2 independent experiments, Mann-Whitney test). (C) Time lapse images of the centrosome in a cell in contact with a droplet (outlined in blue). Distance over time between the centrosome and droplet surface for BSA- or *α*IgG-coated droplets (Mean±SEM). Average plateau distance (25-30 min), (Median±IQR, BSA N=13, *α*IgG N=25, 2 independent experiments, Mann-Whitney test). (D) Time lapse images of the Golgi body in a cell in contact with a droplet (outlined in blue). Distance over time between the Golgi body and droplet surface for BSA- or *α*IgG-coated droplets (Mean±SEM). Average plateau distance (25-30min), (Median±IQR, BSA N=12, *α*IgG N=19, 2 independent experiments, Mann-Whitney test). (E) Time lapse images of lysosomes in a cell in contact with a droplet (outlined in blue). Average distance over time between lysosomes and droplet surface for BSA- or *α*IgG-coated droplets (Mean±SEM). Minimum distance (3-10min), (Median±IQR, BSA N=19, *α*IgG N=32, 2 independent experiments, Mann-Whitney test. (F) Time lapse images of the nucleus in a cell in contact with a droplet (outlined in blue). Nucleus-droplet distance in time (Mean±SEM). Average distance in the final state (25-30 min), (Median±IQR, BSA N=23, *α*IgG N=34, 2 independent experiments, Mann-Whitney test).(G) Characteristic times of polarization events, extracted from the data of (A)-(F) and Figure 3. N*_DAG_*=20, N*_Actin_*=26, N*_Centrosome_* =19, N*_Golgi_* =9, N*_Lyso_*=18, N_*Nuc angle*_=22,N*_Nuc transport_* =34. Timelapse: Scale bar 5 μm.

Finally, we observed that the nucleus was transported to the rear of the cell at later time-points (Figure 2F). Closer observation revealed that this organelle displayed a biphasic movement: a rotation reoriented the nucleus until its stereotypical lymphocyte nuclear invagination faces the immune synapse (*θ_N_*<45° after ~8 minutes); once the nucleus had reoriented, it started moving towards the cell rear ~15 minutes after contact with the droplet, slowly reaching the opposite cell pole over time (Figure 3A-D, Figure 2G).

**Fig. 3.**
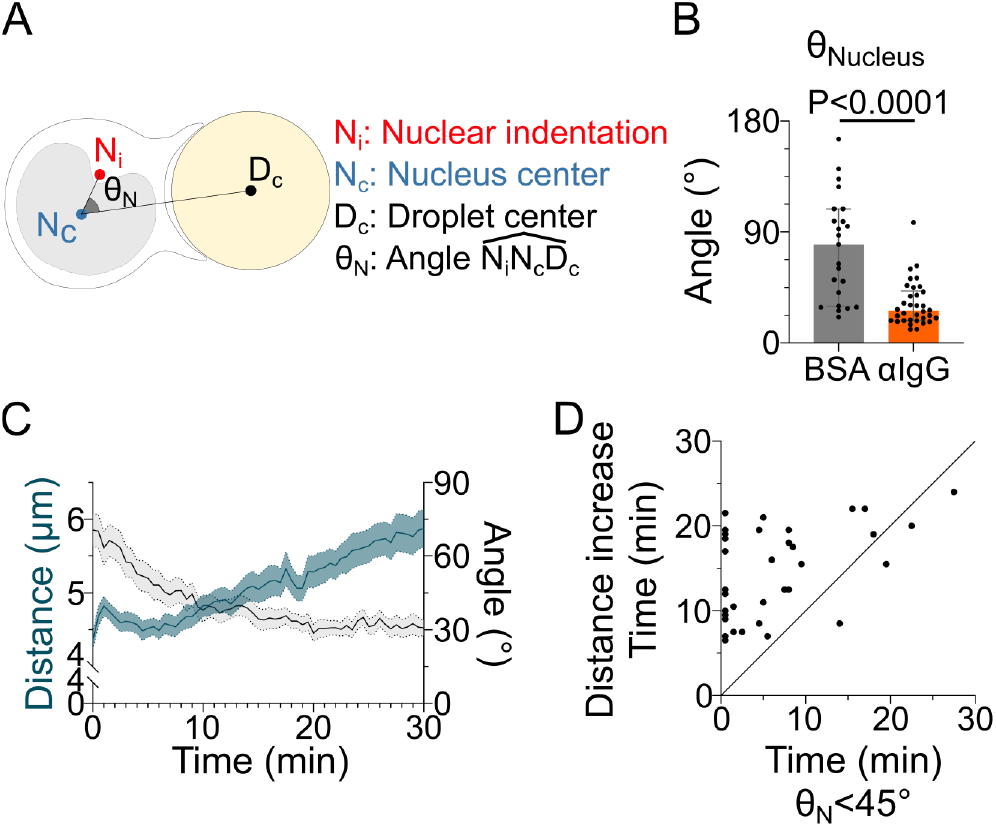
The nucleus undergoes a rotation followed by rearward transport. (A) Schematic defining the angle measured to assess nucleus orientation (Analysis was done in 3D). The indentation was detected based on local curvature. (B) Average angle *θ_N_* in the final state (25-30 min) (Median±IQR, BSA N=23, *α*IgG N=34, 2 independent experiments, Mann-Whitney test). (C) Overlay of nucleus-droplet distance and *θ_N_* over time for cells in contact with *α*IgG-coated droplets and (D) time for which the cell reaches *θ_N_* < 45° (invagination oriented towards the immune synapse), and time of last local minima of nucleus-droplet distance (time after which the nucleus is only transported to the rear) (N=34, 2 independent experiments). Line at Y=X.

In summary, quantification from single kinetics of the various events leading to immune synapse formation in B lymphocytes suggests the existence of two groups of processes: (1) ”early processes” localized at the immune synapse, such as the strong polymerization of F-actin, antigen clustering, and signaling downstream of BCR engagement, which take place in the first 3 minutes; (2) global rearrangements resulting in the reorientation of the centrosome, Golgi apparatus and nuclear invagination to the immune synapse, the recruitment of lysosomes, and later on, the rearward transport of the nucleus. These local and global events associated to synapse formation will be referred to as early and late events from now on.

### The actin cytoskeleton is needed for early but not late events of synapse formation

Having identified the temporal sequence of trafficking events associated to immune synapse formation, we next investigated their interdependency and coordination by the actin and microtubule cytoskeletons. We found that inhibition of actin polymerization with Latrunculin A drastically impaired the clustering of antigen at the droplet surface (Figure 4A), as well as the production of DAG downstream of BCR signaling (Figure 4B). However, neither inhibition nor activation of Myosin II contractility (using the inhibitor para-nitroBlebbistatin or the TRPML1 Calcium channel agonist MLSA1 (12, 25)) strongly affected antigen clustering and DAG production (Figure Supp 1A, B). Taken together, these results stress the importance of F-actin organization -but not actomyosin contractility-in early local events of immune synapse formation, namely antigen clustering and BCR signaling.

**Fig. 4.**
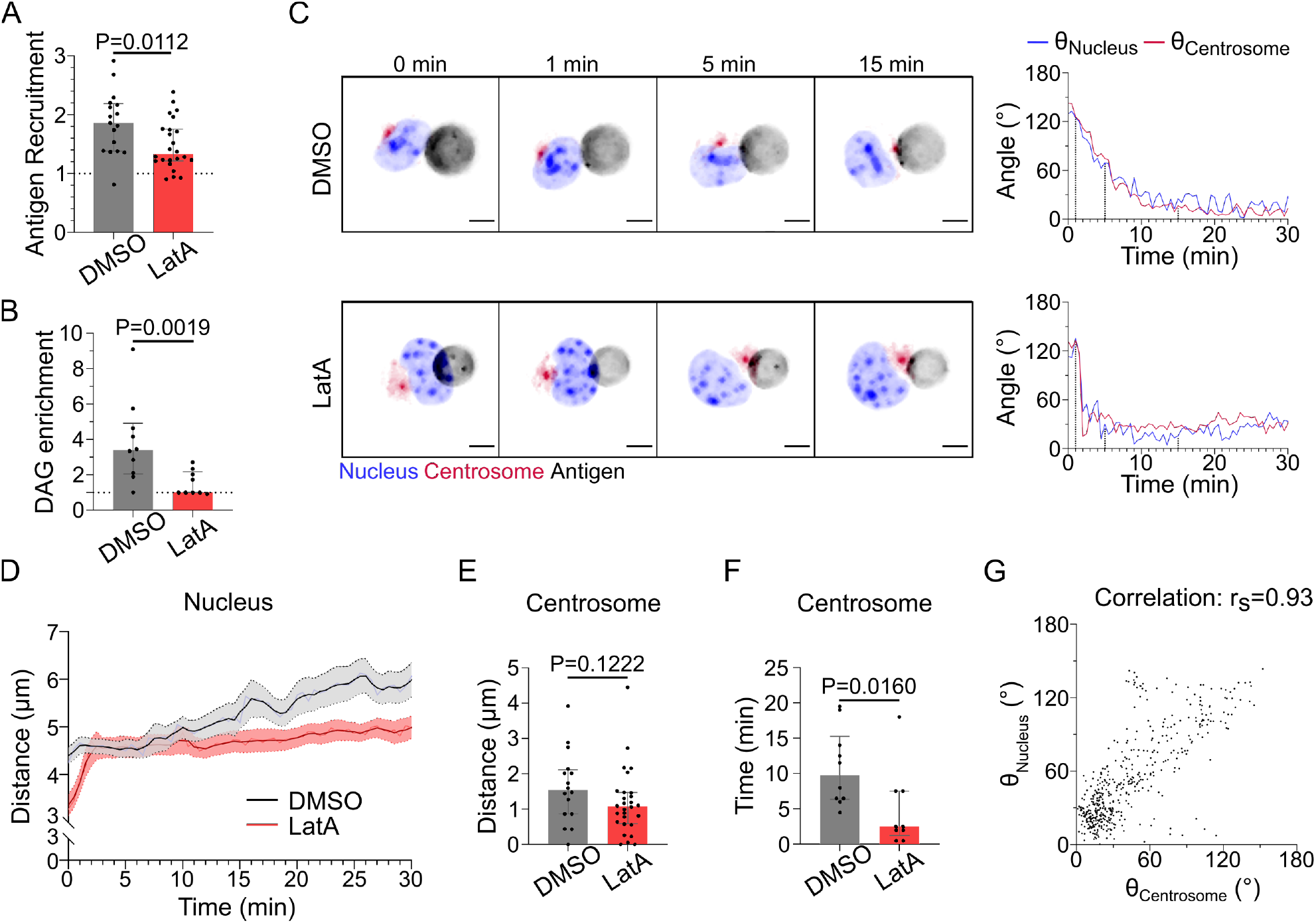
F-actin is essential for antigen recruitment and signaling amplification, but not for the establishment of the polarity axis. (A) Plateau of antigen recruitment (average values 25-30min). Line at Antigen recruitment=1(uniform fluorescence on the droplet). Median±IQR, DMSO N=17, LatA N=24, 2 independent experiments, Mann-Whitney test. (B) Maximum DAG enrichment (in 0-10min). Median±IQR, DMSO N=10, LatA N=9, 2 independent experiments, Mann-Whitney test. (C) Time lapse images of untreated (DMSO) or LatA-treated cells, centrosome in red, nucleus in blue, antigen in gray. Scale bar 5 μm. Right: Angle between the cell-droplet axis and the cell-nucleus invagination (blue) or cell-centrosome (red) axis in time. (D) Nucleus-droplet distance over time. Mean±SEM, DMSO N=17, LatA N=32, 2 independent experiments. (E) Average centrosome-droplet distance (25-30 min). Median±IQR, DMSO N=16, LatA N=28, 2 independent experiments, Mann-Whitney test. (F) Time of centrosome polarization (threshold distance<2 μm). Median±IQR, DMSO N=10, LatA N=9, 2 independent experiments, Mann-Whitney test. (G) Nucleus orientation and centrosome orientation (defined as in Figure 3(A)) during the first 15 min, for DMSO-treated cells. N=16 cells, 1 image every 30 s, 2 independent experiments. Nonparametric Spearman correlation between nucleus-centrosome pairs of data, average correlation 0.93, Confidence interval: 0.86 to 0.97.

Interestingly, imaging centrosome and nucleus re-positioning to the synapse revealed that in the absence of F-actin, these global polarization processes were preserved, and did even take place faster (Figure 4C-F, Supp Movie 4). This acceleration in centrosome polarization might result from loss of F-actin-dependent tethering of this organelle to the nucleus in Latrunculin A-treated cells. Indeed, we previously showed that this pool of F-actin must be cleared for the centrosome to move towards the immune synapse (19). We observed that the centrosome faces the nuclear invagination throughout immune synapse formation, and that they reorient together to ultimately face the immune synapse independently of F-actin (Figure 4C). This was confirmed by the strong correlation between centrosome and nucleus orientation with respect to the cell-droplet axis (Figure 4G). These findings suggest that the centrosome and the nucleus reorient together, which is not affected by F-actin depolymerization.

We conclude that the actin cytoskeleton is essential for the local, early events (Antigen clustering and DAG production downstream of BCR signaling) of synapse formation, but not for the global, late ones (centrosome and nucleus polarization).

### The microtubule cytoskeleton controls both local and global events of synapse formation

Having established how F-actin impacts immune synapse formation, we next addressed its dependency on the microtubule cytoskeleton. For this, we treated cells with Nocodazole to depolymerize microtubules. As expected, microtubule depolymerization prevented centrosome polarization (Figure 5A). Nucleus polarization was also impaired (Figure 5B). These findings are consistent with these two organelles re-positioning together, as described above, and further suggest that their movement is driven by microtubules.

**Fig. 5.**
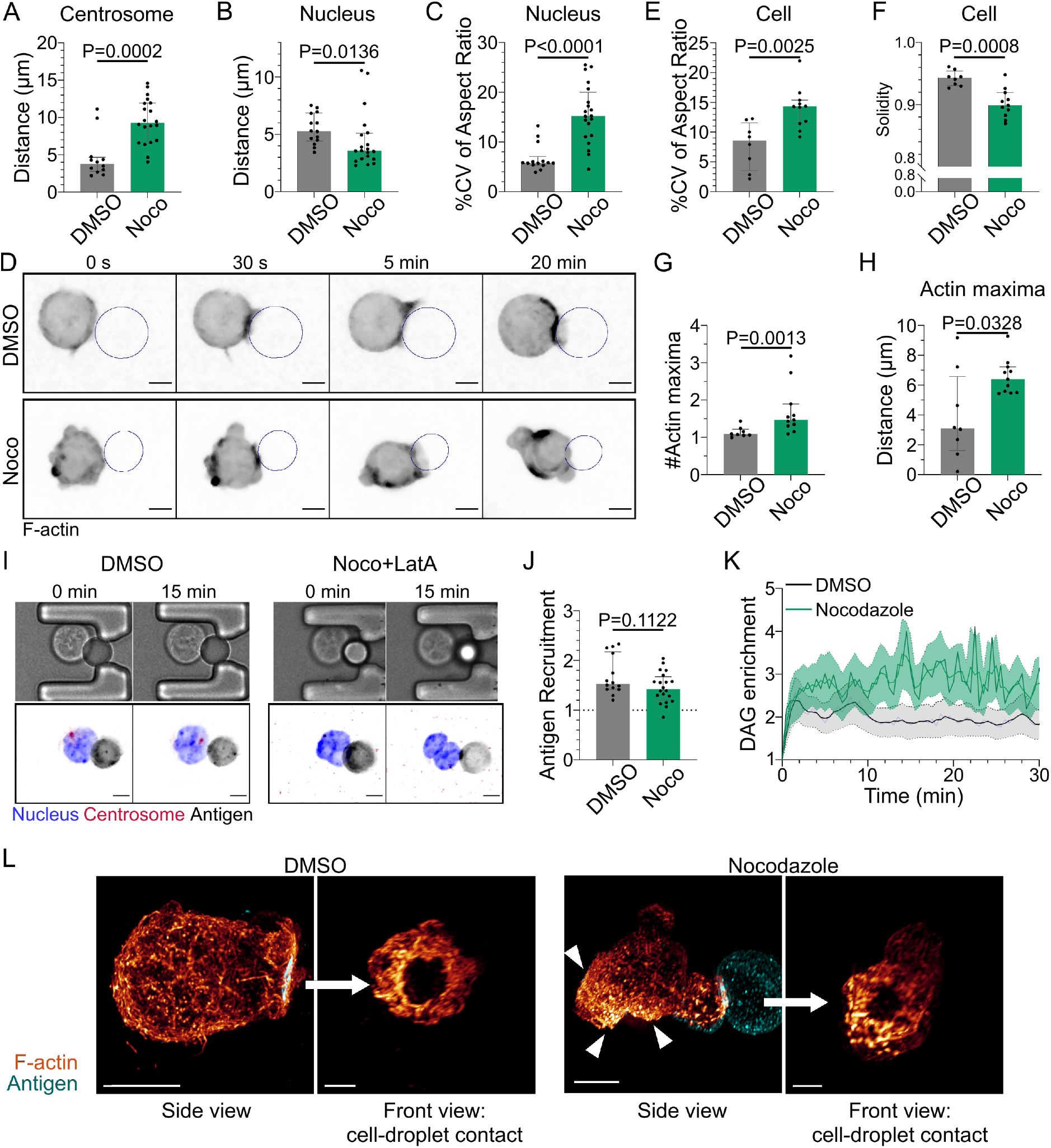
Microtubule disruption leads to intense cell and nucleus deformation, and impairs the establishment and maintenance of a polarized organization. (A) Average centrosome-droplet distance (25-30 min) (Median±IQR, DMSO N=12, Noco N=20, 2 independent experiments, Mann-Whitney test). (B) Average Nucleus-droplet distance (25-30 min) and (C) %Coefficient of Variation of 2D aspect ratio of individual nuclei over time (Median±IQR, DMSO N=14, Noco N=20, 2 independent experiments, Mann-Whitney test). (D) Time lapse images of F-actin in DMSO- or Nocodazole-treated cells, droplet outlined in blue. Scale bar 5 μm. (E) %Coefficient of Variation of 2D aspect ratio of individual cells over time and (F) Median 2D solidity of individual cells (Median±IQR, DMSO N=8, Noco N=11, 2 independent experiments, Mann-Whitney test). (G) Average number of F-actin maxima detected per cell over time and (H) Average distance of maxima to the droplet surface (Median±IQR, DMSO N=8, Noco N=11, 2 independent experiments, Mann-Whitney test). (I) Example images of untreated (DMSO) or treated (Noco+LatA) cells, Bright Field and Fluorescence (eGFP-Cent1, Hoechst, Antigen). Scale bar 5 μm. (J) Plateau of antigen recruitment on the droplet (average values 25-30 min) (Median±IQR, DMSO N=14, Noco N=20, 2 independent experiments, Mann-Whitney test). (K) DAG enrichment over time (Mean±SEM, DMSO N=13, Noco N=10, 2 independent experiments). (L) Examples of 3D SIM immunofluorescence imaging of F-actin and antigen on the droplet after 15-20 minutes of immune synapse formation. White arrowheads: sites of actin enrichment outside of the immune synapse. Side view: Scale bar 5 μm. Front view: Scale bar 2 μm.

Remarkably, we observed that microtubule depolymerization induced major events of nucleus and cell deformation (Figure 5C-E, Supp Movies 5,6) as well as blebbing (Figure 5F). These deformation events were associated to aberrant F-actin distribution: multiple F-actin polymerization spots were visible all around the cell, even far from the immune synapse (Figure 5D, G, H). Accordingly, depolymerizing F-actin in Nocodazole-treated cells with Latrunculin A restored their round shape (Figure 5I). Microtubule depolymerization had a mild impact on antigen clustering and DAG signaling (clustering was slightly reduced while DAG was slightly more sustained) (Figure 5J, K). In addition, morphological analysis of the synapse showed that the stereotypical concentric actin patterning at the immune synapse was preserved (Figure 5L).

Altogether, these results show that microtubules are instrumental for the global late events of synapse formation (centrosome and nucleus re-positioning), but also suggest that microtubules maintain the polarization axis of the cell by limiting the polymerization of the actin cytoskeleton to the immune synapse, consistent with a role for these filaments in synapse maintenance.

### Microtubules restrict actin polymerization to the immune synapse via GEF-H1 and RhoA

How do microtubules restrict actin polymerization to allow its accumulation at the immune synapse and prevent aberrant non-polarized actin distribution? A good candidate to be involved in this process is the guanine exchange factor H1 (GEF-H1), an activator of the RhoA small GTPase that is released from microtubules upon depolymerization (26). GEF-H1 was recently shown to be activated at the immune synapse upon microtubule acetylation in activated B lymphocytes, which could help them tuning RhoA activity to locally control actin dynamics (27).

To test the involvement of GEF-H1 in microtubule-dependent regulation of cortical actin distribution, we silenced its expression using siRNA (Figure 6A). Strikingly, GEF-H1 silencing prevented cell deformation and blebbing upon microtubule depolymerization (Figure 6B-D) and restored the polarized polymerization of actin at the immune synapse (Figure 6E). These results indicate that the aberrant non-polarized actin polymerization observed upon treatment of B lymphocytes with Nocodazole most likely results from GEF-H1 release from microtubules. Accordingly, we found that B cells expressing a constitutive active form of RhoA (RhoA L63, referred to as RhoA CA) displayed a phenotype similar to the one of Nocodazole-treated cells: aberrant actin polymerization, dynamic cell deformation and blebbing (Figure 6F-I, Supp Movie 7).

**Fig. 6.**
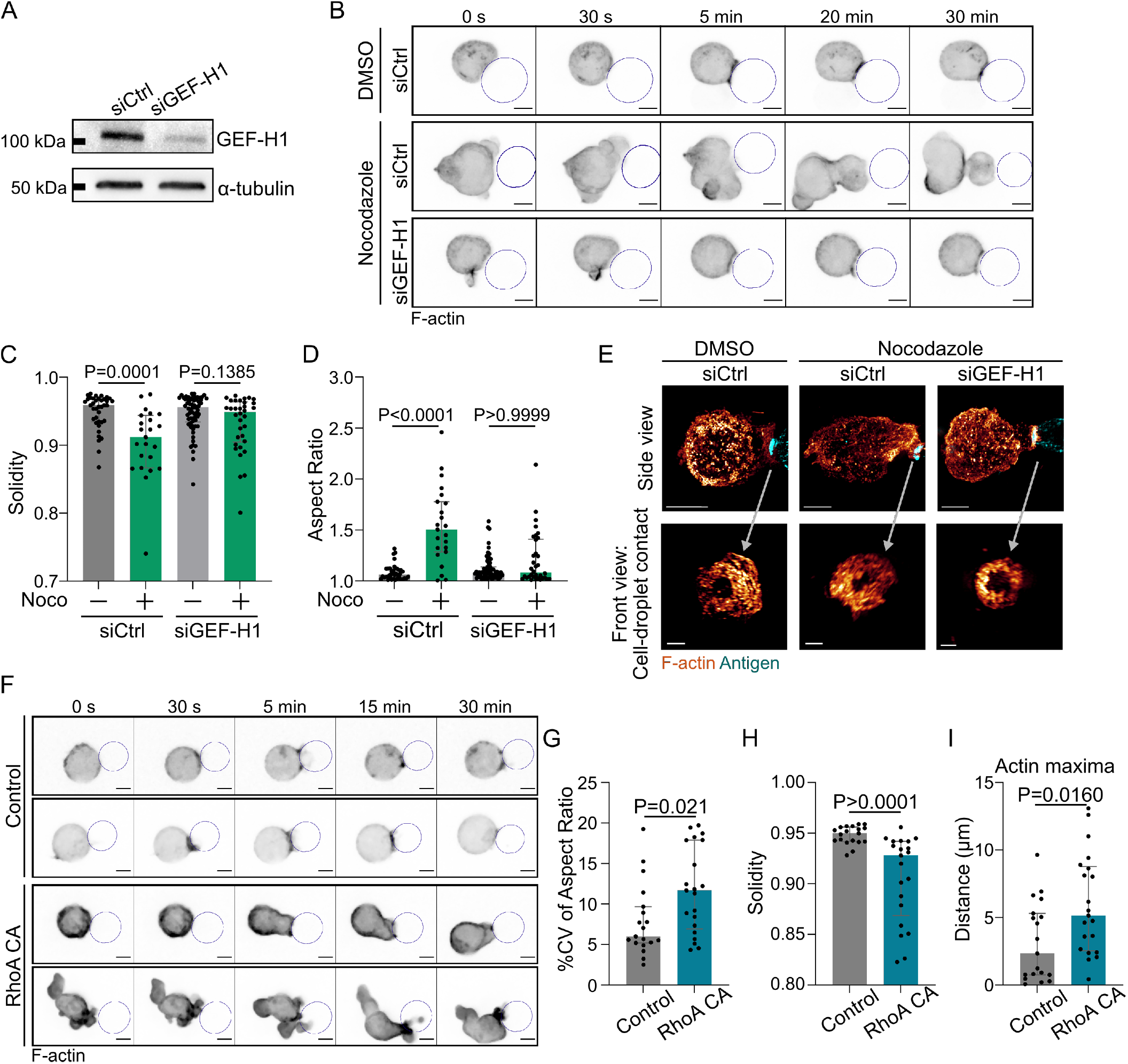
GEF-H1/RhoA is responsible for cell shape and actin patterning defects upon microtubule depletion. (A) Western blot quantification of the efficiency of GEF-H1 silencing. *α*-tubulin was used as a loading control. The blot presented is representative of 2 independent experiments. (B) Time lapse images of F-actin in cells transfected with siCtrl or siGEF-H1 and treated with DMSO (control) or Nocodazole. Scale bar 5 μm. (C) Solidity in 2D and (D) Aspect ratio of cells after 40min of immune synapse formation (siCtrl DMSO N=38, siCtrl Noco N=23, siGEF-H1 DMSO N=65, siGEF-H1 Noco N=34, 2 independent experiments, Kruskal-Wallis test with multiple comparisons between DMSO and Noco). (E) Examples of 3D SIM immunofluorescence imaging of F-actin and antigen on the droplet after 15-20 min of immune synapse formation. Side view: Scale bar 5 μm. Front view: Scale bar 2 μm. (F) Time lapse images of F-actin in cells transfected with a control empty vector (pRK5) or expressing RhoA CA (constitutively active). Scale bar 5 μm. (G) %Coefficient of Variation of 2D aspect ratio of individual cells over time, (H) Median 2D solidity of individual cells and (I) Average distance of actin maxima to the droplet surface (Median±IQR, Control N=19, RhoA CA N=21, 2 independent experiments, Mann-Whitney test).

The activation of RhoA by GEF-H1 leads to both nucleation of linear actin filaments by diaphanous formins (mDia) and activation of myosin II by the ROCK kinase for contraction of these filaments (28, 29). We, therefore, asked whether modulation of actin nucleation or myosin II activity had any impact on the phenotype of Nocodazole-treated cells. Noticeably, we found that while Myosin II inhibition (using para-nitroBlebbistatin) prevented cell blebbing upon microtubule depolymerization (Figure 7A, B), it did not restore cell shape, with cells elongating over time (Figure 7A, C, D), nor polarized actin polymerization (Figure 7A, E, Supp Movie 8). These results suggest that actin nucleation, rather than myosin II activation, downstream of GEF-H1 and RhoA activation is responsible for the non-polarized polymerization of actin upon microtubule depolymerization. Accordingly, simultaneous depolymerization of actin and microtubules prevented cell deformation, restoring both cell and nucleus shape (Figure 5I).

**Fig. 7.**
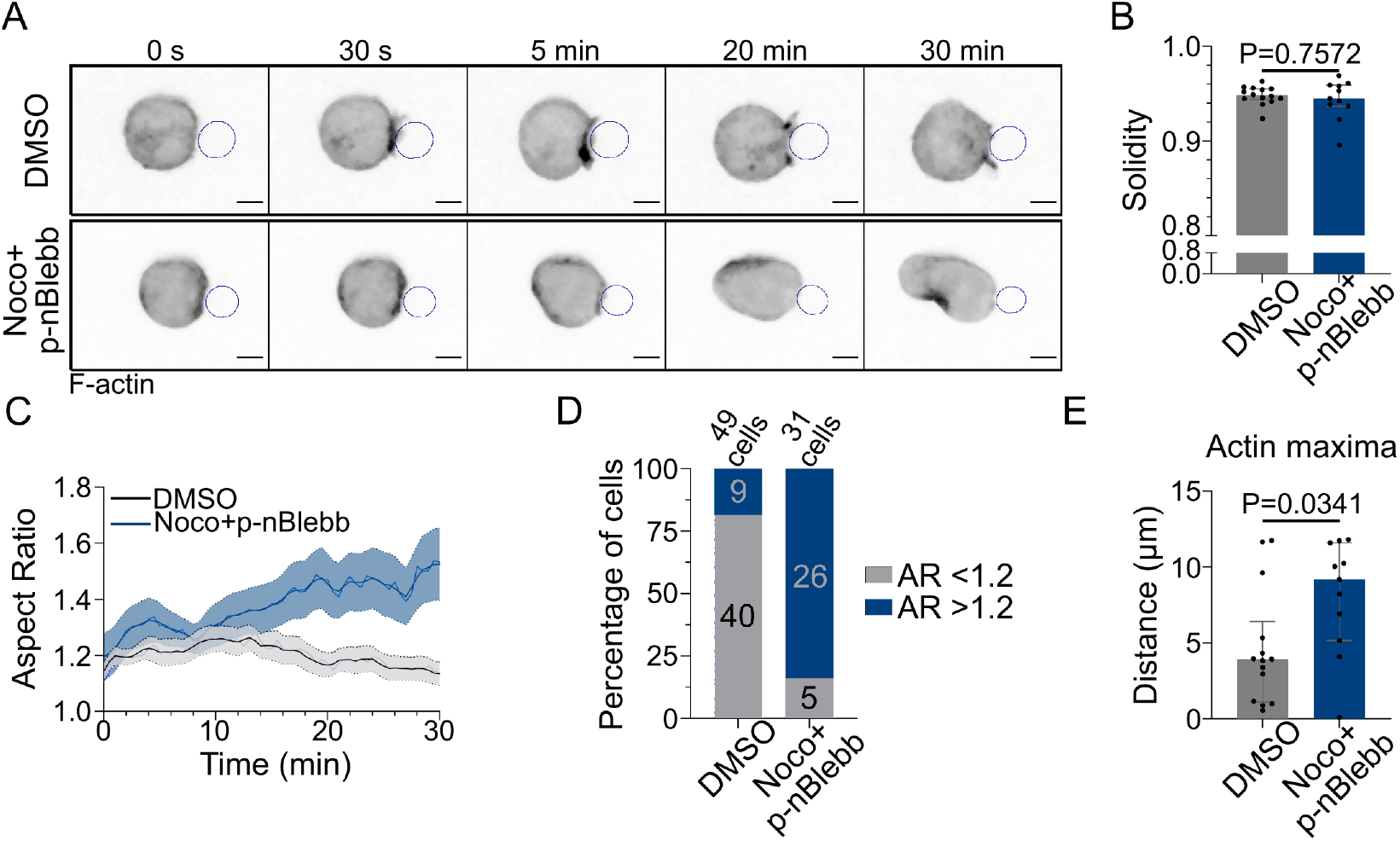
Regulation of polarized actin polymerization by microtubules is Myosin II-independent. (A) Time lapse images of F-actin, droplet outlined in blue. Scale bar 5 μm. (B) Median 2D solidity of individual cells over time (Median±IQR, DMSO N=14, Noco+p-nBlebb N=11, 3 independent experiments, Mann-Whitney test). (C) Aspect ratio of cells in time (Mean±SEM, DMSO N=14, Noco+p-nBlebb N=11, 3 independent experiments). (D) Percentage of cells with Aspect Ratio >1.2 or <1.2 after 40 min of synapse formation. (E) Average distance of F-actin maxima to the droplet over 30 min of synapse formation (Median±IQR, DMSO N=14, Noco+p-nBlebb N=11, 3 independent experiments, Mann-Whitney test).

### Restriction of actin nucleation by microtubules promotes the formation of a unique immune synapse

Our results suggest that by titrating GEF-H1, microtubules tune the level of RhoA activation to restrict actin polymerization to the immune synapse, thus stabilizing a single actin polarity axis. We hypothesized that such regulatory mechanism might help B cells maintaining a unique immune synapse, rather than forming multiple synapses all over their cell body. To test this hypothesis, we first allowed an immune synapse to form between a cell and a droplet, then injected a second round of droplets as soon as possible (<3 minutes after the first contact), allowing the cell to contact several antigen-coated droplets. For this experiment, we chose to use cells treated with both Nocodazole and para-nitroBlebbistatin to prevent excessive blebbing and facilitate the analysis. We observed two types of cell behaviors: they either brought the droplets together into a single immune synapse, or formed multiple, separated immune synapses (Figure 8A, B). Noticeably, microtubule-depleted cells formed more multiple separated synapses than control cells (Figure 8C). Accordingly, while control cells were able to merge contacted droplets into a unique immune synapse, this was not observed in cells whose microtubule were depolymerized (Figure 8B, Supp Movie 9). These results are consistent with microtubule being required for the formation and maintenance of a unique immune synapse, wherein F-actin polymerization concentrates, rather than multiple dispersed ones. To test this hypothesis, we computed the difference between the brightest and the dimmest synapse in terms of F-actin enrichment. We found that in control cells binding multiple droplets, F-actin was often enriched at one single synapse (Figure 8D). By contrast, microtubule-depleted cells exhibiting multiple synapses rather showed equal F-actin distribution from one synapse to another. These results strongly suggest that microtubule depolymerization leads to competition of nascent synapses for F-actin, favoring the formation of multiple synapses rather than a unique one. We therefore conclude that, by restricting RhoA-dependent actin polymerization via GEF-H1, microtubules allow the maintenance of a single polarity axis and stabilizes in space and time the immune synapse of B lymphocytes.

**Fig. 8.**
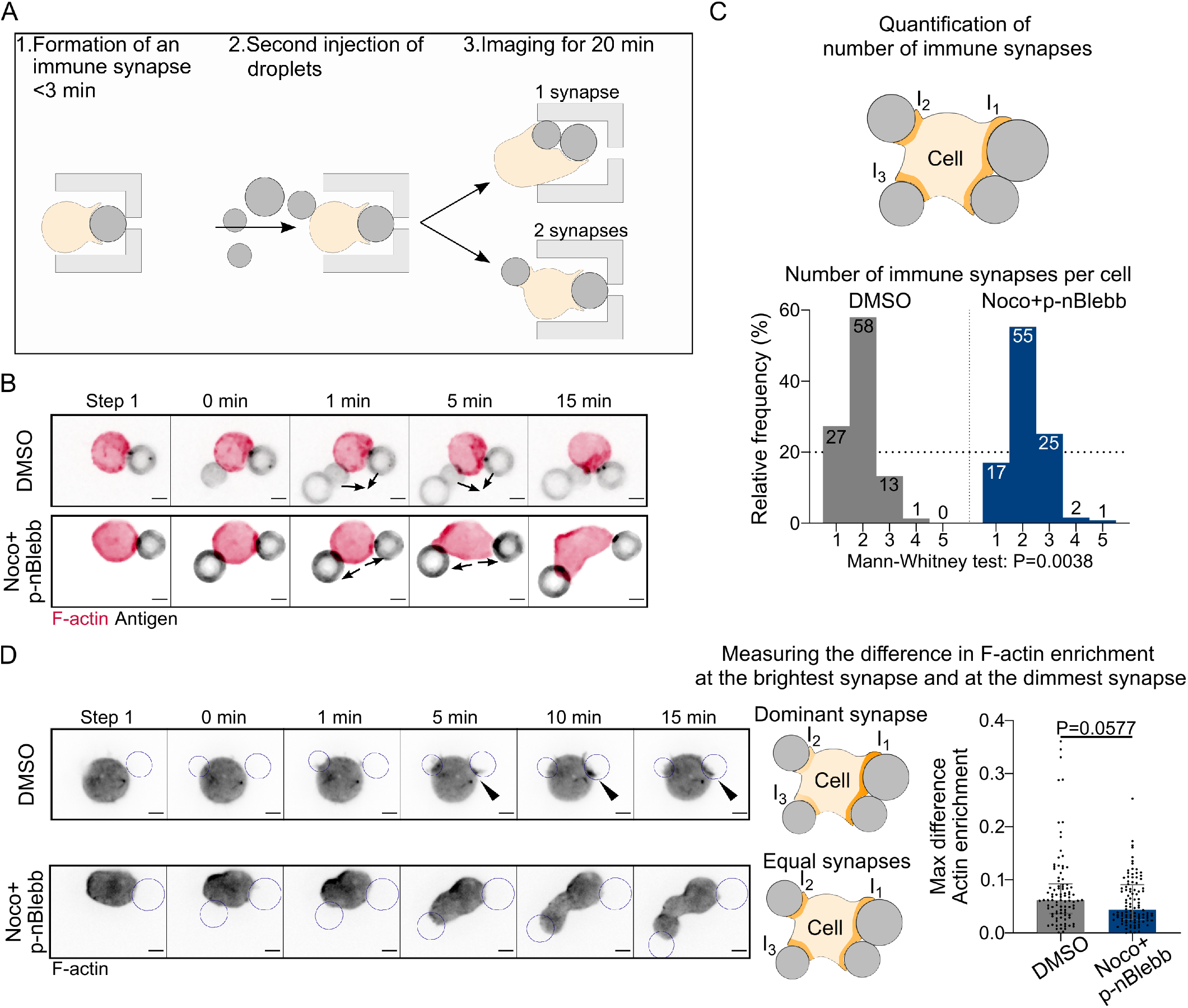
Microtubule depletion favors the formation of multiple immune synapses. (A) Schematic of the concept of the “sandwich” experiment. (B) Examples of time lapse images of F-actin and antigen on the droplet. Situation of a cell (untreated) bringing droplets closer into one immune synapse, and of a cell (treated with Nocodazole and para-nitroBlebbistatin) taking droplets apart. Scale bar 5 μm. (C) Number of immune synapses per cell after 20 minutes of interaction (DMSO N=150, Noco+p-nBlebb N=123, 2 independent experiments, Mann Whitney test P=0.0038). (D) (Left) Examples of time lapse images of F-actin and antigen on the droplet. Situation of a cell (untreated) with one synapse more enriched in F-actin, and of a cell (treated with Nocodazole + para-nitroBlebbistatin) with equivalent synapses. Scale bar 5 μm. (Right) Quantification of the difference of enrichment in F-actin at the brightest and the dimmest immune synapse, per cell (DMSO N= 94, Noco+p-nBlebb N=104, 2 independent experiments, Mann-Whitney test).

## Discussion

In this work, we used a custom microfluidic system to study the coordination by actin and microtubule cytoskeletons of the various events associated to immune synapse formation in B lymphocytes. We observed that this process is characterized by two classes of events: a first phase (in the first 3.5 minutes), where F-actin is strongly polymerized at the site of contact, leading to antigen accumulation and production of DAG as a result of BCR signaling, and a second phase during which the centrosome is reoriented towards the immune synapse together with the Golgi apparatus and lysosomes while the nucleus undergoes a rotation followed by backward transport. We found that F-actin polymerization is only needed for the first phase, in contrast to microtubules that not only control centrosome and organelle re-positioning but further maintain a unique polarity axis by restricting actin nucleation to the immune synapse. We propose that this mechanism might allow reinforcing a single synapse rather than multiple competing ones.

How do microtubules restrict F-actin polymerization to the immune synapse? We identified GEF-H1 as a key player in this process, which limits RhoA activity and downstream actin nucleation to the synapse. Indeed, we observed that global activation of GEF-H1-RhoA axis induced actin polymerization outside of the synapse, independently of myosin II activity. Interestingly, it was recently shown that microtubules were acetylated in the vicinity of the centrosome upon immune synapse formation, resulting in the local release and activation of GEF-H1 (27, 30). Our results suggest that GEF-H1 might activate RhoA to trigger downstream formin-dependent actin nucleation at the immune synapse exclusively. In this model, RhoA would remain inactive in the rest of the cell, most likely due to GEF-H1 trapping on microtubules deacetylated by HDAC6 (30, 31). We suggest that this “local activation” of GEF-H1 and “global inhibition” by trapping is a reminiscent of the Local Excitation Global Inhibition model, described in amoeba, where symmetry breaking arises from a local positive feedback (PIP3 that promotes F-actin polymerization) combined to a globally active diffusible inhibitory signal (PTEN, a PIP3 phosphatase) (32–34).

The Local Excitation Global Inhibition model predicts the establishment of a single stable polarity axis. Accordingly, our experiments show that unperturbed B lymphocytes favor the formation of a unique synapse over multiple ones, even when particulate antigens are presented from several locations. We propose that this mechanism, at least in enzymatic extraction, could help improving antigen extraction. Indeed, GEF-H1 has been shown to be necessary for the assembly of the exocyst complex at the immune synapse, and therefore for protease secretion (27). In this context, the localized release and activation of GEF-H1 by microtubules at the immune synapse might allow for the concentration of resources, promoting F-actin polymerization and optimizing proteolytic extraction at one unique site. Polarization of the centrosome and reorientation of the microtubule network would thus reduce the dispersion of resources in secondary synapses. Indeed, the release of proteases in several locations, or in an open environment (as opposed to the tight synaptic cleft) could result in a lower local concentration of proteases, and therefore lower the efficiency of antigen uptake. A unique polarity axis could also be beneficial to T/B cooperation as upon antigen internalization, B cells must migrate to the T cell zone for antigen presentation, the speed and persistence of migration being directly related to the robustness of cell polarity (4, 35). In addition, it has been shown that B cells can undergo asymmetric cell division upon synapse formation and antigen extraction, which prevents antigen dilution upon cell division, an event that also require a stable polarity axis (36, 37). Future experiments aimed at studying how these downstream events of synapse formation are regulated when B cells nucleate actin all over their cell cortex and form multiple contacts should help address these questions.

In conclusion, we showed that microtubules can act as master regulators of actin nucleation, maintaining the formation of a single immune synapse in B lymphocytes. This control relies on the GEF-H1-RhoA axis, which may be at the core of a “Local Excitation Global Inhibition” model. Our work points at the interaction between actin and microtubules as a way to control the axis of cell polarity, that might be common to a larger class of cells.

## Materials and Methods

### Cells and cell culture

The mouse IgG^+^ B lymphoma cell line IIA1.6 (derived from the A20 cell line [ATCC #: TIB-208]) was cultured as previously reported (14) in CLICK Medium (RPMI 1640 – GlutaMax-I + 10% fetal calf serum, 1% penicillin– streptomycin, 0.1% *β* mercaptoethanol, and 2% sodium pyruvate). Fetal calf serum was decomplemented for 40 min at 56°C. All cell culture products were purchased from GIBCO/Life Technologies. All experiments were conducted in CLICK + 25mM HEPES (15630080, Gibco).

### Antibodies and Reagents

#### For droplet preparation fabrication and functionalization

DSPE-PEG(2000) Biotin in chloroform (Avanti Lipids, Coger 880129C-10mg), Soybean oil (Sigma-Aldrich, CAS no. 8001-22-7), Pluronic F68 (Sigma-Aldrich, CAS no. 9003-11-6), Sodium Alginate (Sigma-Aldrich, CAS no. 9005-38-3), Tween 20 (Sigma Aldrich, CAS 9005-64-5), Na_2_HPO_4_· 7H_2_O (Sodium phosphate dibasic heptahydrate, M=268g/mol, CAS 7782-85-6, Merck), NaH_2_PO_4_ · H_2_O (Sodium phosphate monobasic monohydrate M = 138g/mol, CAS 10049-21-5, Carlo Erba), Streptavidin Alexa Fluor 488 (Thermofisher, S11223), Streptavidin Alexa Fluor 546 (Thermofisher S11225), Streptavidin Alexa Fluor 647 (Thermofisher S32357), biotin-SP-conjugated AffiniPure F(ab′)_2_ Fragment Gt anti Ms IgG (Jackson ImmunoResearch 115-066-072), Biotin labeled bovine albumin (Sigma-Aldrich A8549-10MG).

#### For microfluidic chips

PDMS-RTV 615 (Neyco RTV6115), Polyvinylpyrrolidone K90 (Sigma 81440, called PVP), Medical tubing, Tygon® ND 100-80 (Saint-Gobain), Stainless Steel Plastic Hub Dispensing Needles 23 GA (Kahnetics KDS2312P), Fluorodish (World Precision instruments FD35).

#### Dyes and plasmids for live cell imaging

Hoechst 33342 (Thermofisher, R37605) kept in solution, Lysotracker Deep Red (Thermofisher, L12492) 50 nM in incubator for 45 min then wash, SirTubulin kit (Spirochrome AG, Tebu-bio SC002) 100 nM SiRTubulin+10 μM verapamil >6 h, eGFP-Centrin1 plasmid used in (19), F-tractin tdTomato obtained from the team of Patricia Bassereau (Institut Curie, Paris), Rab6-mCherry plasmid obtained from Stéphanie Miserey (Institut Curie, Paris), C1*δ*-GFP plasmid was obtained from Sergio Grinstein (23). pRK5myc RhoA L63 (RhoA CA - constitutively active) was a gift from Alan Hall (Addgene plasmid 15900; http://n2t.net/addgene:15900; RRID:Addgene_15900) (38), and an empty pRK5myc vector was used as a negative control. Expression of Ftractin-tdTomato, Rab6-mCherry, *C*1*δ*-GFP, pRK5myc and RhoA L63 was achieved by electroporating 1.10^6^ B lymphoma cells with 0.25 to 0.5 μg of plasmid using the 10 μL Neon Transfection system (Thermofisher). Expression of eGFP-Centrin1 was achieved by electroporating 4.10^6^ B lymphoma cells with 4 μg of plasmid using the Amaxa Cell Line Nucleofector Kit R (T-016 program, Lonza). Cells were cultured in CLICK medium for 5 to 16 h before imaging.

For siRNA silencing, IIA1.6 cells were transfected 60-70 h before live experiment with 40 pmol siRNA per 10^6^ cells using the 10 μL Neon Transfection system (Thermofisher) and ON-TARGETplus Control n=Non-Targeting Pool (Dharmacon, D-001810-10-05) or SMARTPool ON-TARGETplus Mouse Arhgef2 siRNA (Dharmacon, L-040120-00-0005).

#### For immunofluorescence and Western Blot

Formaldehyde 16% in aqueous solution (Euromedex, 15710), BSA (Euromedex, 04-100-812-C), PBS (Gibco, 10010002), Rabbit anti GEF-H1 (Abcam, ab155785, 1/1000 for WB), Rat anti *α*-tubulin (Biorad, MCA77G, 1/1000 for WB), Anti-Rabbit IgG, HRP-linked Antibody (Cell signaling, #7074, 1/5000 for WB), Anti-Rat IgG, HRP-linked Antibody (Cell signaling, #7077, 1/10000 for WB), Alexa Fluor 546 Phalloidin (Thermofisher, A22283, 1/200), DAPI (BD Bioscience, 564907, 1/1000), Saponin (Sigma, 8047-15-2), Fluoromount-G (Southern Biotech, 0100-01), RIPA Lysis and Extraction Buffer (Thermofisher, 89900), Protease inhibitor cocktail (Roche, 11697498001), Benzonase (Sigma, E1014-5KU), Laemmli sample buffer (Biorad, 1610747), NuPAGETM Sample reducing agent (Invitrogen, NP0004), Gels, and materials for gel migration and membrane transfer were purchased from Biorad, Clarity™ Western ECL Substrate (Biorad, 1705060).

#### Drugs and inhibitors

Latrunculin A (Abcam, ab144290, incubation 2μM for 1 h), para-nitroBlebbistatin (Optopharma, 1621326-32-6, incubation 20 μM for 1 h), Nocodazole (Sigma, M1404, incubation 5μM for 1 h), MLSA1 (Tocris, 4746, incubation 1 μM for 1 h). For all experiments in microfluidic chips involving drugs, chips were filled with media+drug (or DMSO) at least 1 h before experiment, and only media+drug was used at each step.

### Experimental protocols

#### Droplet stock formulation

Oil phase: 150 μL of DSPE-PEG(2000) Biotin solution (10 mg/mL in chloroform) in 30 g of soybean oil, left >4 h in a vacuum chamber to allow chloroform evaporation. Aqueous phase: 10 g of 1% Sodium alginate, 15% Pluronic F68 solution in deionized water, gently mixed with a spatula to avoid bubbles. The oil phase was slowly added to the aqueous phase, starting by 2-3 drops, gentle stirring until oil was incorporated, then repeating. Over time, the oil phase incorporates more easily and could be added faster, until a white emulsion was obtained. The emulsion was then sheared in a Couette cell (39) at 150 rpm to obtain droplets of smaller and more homogeneous diameter. The new emulsion was recovered as it got out of the Couette cell, and was now composed of 25%^v^/v aqueous phase containing 15%^w^/v Pluronic F68. To wash and remove the smallest droplets, the droplet emulsion was put in a separating funnel for 24 h at 1% Pluronic F68, 5% oil phase. This operation was repeated at least 2 times. The final emulsion was stored in glass vials at 12°C, and droplets had a median diameter of 9.4 μm.

This type of droplets was previously characterized using the pendant drop technique (40, 41), and appear like a relatively stiff substrate (surface tension 12 mN.m^−1^ measured by the pendant drop technique (42), equivalent to a Laplace pressure of 4.8 kPa for a droplet of radius 5 μm). The antigen concentration is estimated to be of the order of 50 mol/μm^2^ (see (43) for method) and the diffusion constant ~0.7 μm^2^ .s^−1^, measured by FRAP, comparable to lipid bilayers (2, 44–46).

#### Droplet functionalization

Droplets were functionalized on the day of experiment. All steps were performed in low binding eppendorfs (Axygen Microtubes MaxyClear Snaplock, 0.60 ml, Axygen MCT-060-L-C), and using PB+Tween20 buffer (Tween 20 at 0.2%^v^/v in PB Buffer pH=7, 20 mM). A small volume of droplet emulsion (here 2 μL) was diluted 100 times in PB+Tween20 buffer, and washed 3 times in this buffer. Washes were performed by centrifugating the solution for 30 s at 3000 rpm in a minifuge, waiting 30 s and then removing 170 μL of the undernatant using a gel tip, then adding 170 μL of PB+Tween20. At the last wash, a solution of 170 μL + 2.5 μL of fluorescent streptavidin solution (1 mg/mL) was added to the droplet solution, then left on a rotating wheel for 15 min, protected from light. Droplets were then washed 3 times, and at the last wash a solution of 170 μL PB+Tween20 + 5 μL of Biotin Goat F(ab′)2 anti-Mouse IgG (1 mg/mL) (or other biotinylated protein in the same proportion) was added and left to incubate for >30 min on a rotating wheel, protected from light. Droplets were finally washed three times before use, with PB+Tween20. For experiments using drug treatments, droplets were resuspended in culture media + drug before the experiment.

#### Microfluidic chip fabrication

Microfluidic chips were made using an original design from the team of Jacques Fattaccioli (ENS Paris, IPGG) (47). RTV PDMS was mixed at a ratio 1:10, and poured in epoxy cast replicates of the microfluidic chips, and cooked until fully polymerized. Microfluidic chips were then cut, and 0.5 mm diameter holes were made at the entry/exit sites. The PDMS chip and a Fluorodish were then activated in a plasma cleaner (PDC-32G Harrick) for 1 min and bonded to each other for 1 h at 60°C. Bonded chips were activated in the plasma cleaner for 1 min to be activated, and filled using a syringe with a 0.2%*^w^/^_v_^* PVP K90 solution in MilliQ water, to form an hydrophilic coating. Microfluidic chips were then kept at 4°C in the 0.2%*^w^/^_v_^* PVP K90-filled fluorodish to prevent drying, for up to a week before the experiment. On the day of the experiment, microfluidic chips were moved gradually to room temperature, then into a incubator, before imaging. For experiments using drug treatments, microfluidic chips were injected with culture media + drug in the morning, and left to incubate to ensure stable drug concentration during the experiment.

#### Live imaging of IIA1.6 cell polarization in microfluidic chips

Live imaging of polarization was performed using an inverted spinning disk confocal microscope (Eclipse Ti Nikon/Roper spinning head) equipped with a Nikon 40x, NA 1.3, Plan Fluor oil immersion objective, a CMOS BSI photometrics camera (pixel size 6.5 μm), and controlled with the Metamorph software (Molecular Device, France). Stacks of 21 images (*δ*z=0.7 μm) were taken every 30 s during 40 min, with a binning of 2. Auto Focus was implemented in Metamorph using the Bright Field image, then applied to fluorescent channels with a z-offset at each time point. On the day of the experiment, droplets were functionalized and cells were resuspended at 1.5.10^6^ cells/mL in CLICK+25 mM HEPES. Microfluidic chips, cells and media were kept in an incubator at 37°C with 5% CO_2_ until imaging.

Droplets (diluted 1/6 from functionalized solution) were injected in the microfluidic chip using a Fluigent MFCS™-EZ pressure controller, Tygon tubing and metal injectors from the dispensing needles 23GA. When enough traps contained a droplet, the inlet was changed to CLICK+25 mM HEPES (or CLICK+25 mM HEPES+drug) to rinse PB+Tween20 buffer and remove any antigen in solution or droplet that could remain. After a few minutes, the inlet was changed to the cell suspension, keeping a minimum pressure to avoid cells encountering droplets before acquisition was launched. Stage positions were selected and the acquisition was launched. After one time point (to have an image of droplets without cells, and ensure to have the first time of contact), the inlet pressure was increased to inject cells and create doublets. After 2-5 min (when enough doublets had formed), the injection pressure was lowered to a minimum to limit cell arrival, and perturbation of cells by strong flows.

For multiple synapse experiments, images were acquired every 1 minute, for 20 minutes.

#### Immunofluorescence with droplets

To approach the non-adherent condition of the cells in the microfluidic chips, IIA1.6 cells were seeded for 15 minutes on glass coverslips (Marienfeld Superior Precision Cover Glasses, 12 mm diameter) coated with 100 μg/mL BSA, on which they display limited spreading. Droplets were prepared as for live imaging, then diluted 13 times in CLICK+HEPES. A small volume of this droplet solution was deposited on parafilm, and the coverslip was then flipped onto the droplets and left for 5 minutes, so that droplets would float up to encounter the cells. Coverslips were then put in pre-heated CLICK+HEPES media in a 12-well plate, with the cells facing up, for 15 minutes. All manipulations and washed were performed very gently, using cut pipet tips to limit cell and droplet detachment. Samples were fixed for 12 min at RT using 4% PFA in PBS, then washed three times with PBS. After 30 min of incubation with PBS/BSA/Saponin 1X/0.2%/0.05%, samples were incubated for 1h at RT with 1/200 Alexa Fluor 546 and 1/1000 DAPI in PBS/BSA/Saponin 1X/0.2%/0.05%, then washed three times with PBS. Samples were then mounted using Fluoromount-G and left at RT until dry.

Samples were imaged by 3D SIM, using a Delta Vision OMX v4 microscope, equipped with an Olympus 100X, NA 1.42, Plan Apo N, oil immersion objective, and EMCCD cameras. Image reconstruction was performed using the SoftWoRx image software, under Linux. 3D visualization for figures were performed using the Imaris Viewer software.

#### Western Blot

B cells were lysed for 10 min at 4°C in RIPA Lysis and Extraction Buffer supplemented with protease inhibitor cocktail, then treated with benzonase. Lysates were spinned for 15 min at 4°C at maximum speed to remove debris, followed by heating of supernatants for 5 min at 95°C with Laemmli sample buffer and NuPAGETM Sample reducing agent. Supernatants were loaded onto gels and transferred to PVDF membranes. Membranes were blocked for 45 min at RT with 5 % BSA in TBS+0.05% Tween20, incubated overnight at 4°C with primary antibodies, then incubated 1 h at RT with secondary antibodies. Membranes were revealed using Clarity™ Western ECL Substrate and chemiluminescence was detected using a BioRad ChemiDoc MP imaging system. Western blots were quantified using ImageLab.

### Image analysis

Image analysis was performed on the Fiji software (48) using custom macros, unless stated otherwise. All codes are available upon request. Single kinetic curves analysis were performed using Rstudio (49). Graphs and statistical analysis were made using GraphPad PRISM version 9.2.0 for Windows, GraphPad Software, San Diego, California USA, www.graphpad.com.

For graphs of polarization in time of BSA vs *α*IgG (Figure 2), a moving average filter of length 3 was applied on the mean and SEM before plotting. The non-smoothed mean curve is superimposed to the graphs.

For image analysis of live imaging, cell-droplet doublets were cropped from original acquisitions, and were cut so that cells arrive at the second frame (marked as 0 s in figures).

#### Analysis of antigen recruitment on the droplet

Bleaching of fluorescent streptavidin was corrected before analysis using Bleach Correction - Histogram Matching. Antigen recruitment was measured by computing the ratio between fluorescence intensity at the synapse and fluorescence intensity at the opposite side on three planes passing through the droplet and the cell, normalized by this value at the time of cell arrival (Figure 1D).

#### Analysis of F-tractin-tdTomato

Fluorescence was corrected using the Bleach Correction-simple ratio program. Using a custom Fiji macro, 3D masks of the droplet and the cell were generated. Enrichment of F-actin at the immune synapse was defined as the sum of intensity in the mask of the cell within a 2 μm layer around the droplet in 3D, divided by the sum of intensity in the mask of the cell. This measurement was normalized by its value at the first time point of encounter between the cell and the droplet to compensate for potential heterogeneity of the initial state. Extraction of characteristic values (time of peak, maximum) were extracted with R, on single kinetic curves smoothed using 3R Tukey smoothing (repeated smoothing until convergence) (50). Time and value of maximum were computed in the first 10 min of cell-droplet contact. Shape characteristics of the cell (aspect ratio, solidity) were measured on maximum z projections of cell masks.

#### Analysis of C1*δ*-GFP DAG reporter

Fluorescence was corrected using the Bleach Correction-simple ratio program. Using a custom Fiji macro, 3D masks of the droplet was generated. Enrichment of C1*δ*-GFP (C1 domain of PKC*δ*, acting as a DAG reporter (23)), was defined as the sum of intensity within a 1 μm layer around the droplet. This measurement was normalized with its value at the first time point of encounter between the cell and the droplet, to account for variability of reporter expression between cells. Extraction of characteristic values (time of peak, maximum, plateau value relative to maximum) were extracted with R, on single kinetic curves smoothed using 3R Tukey smoothing (repeated smoothing until convergence) (50). Time and value of maximum were computed in the first 10 min of cell-droplet contact.

#### Analysis of the centrosome

The 3D movie was first interpolated to obtain isotropic voxels for the advanced analysis. Using a custom Fiji macro, a mask of the droplet was generated, and position of the centrosome (stained with SiRtubulin) was detected, to measure the distance of the centrosome from the droplet surface. Characteristic times were extracted on single kinetic curves smoothed using 3R Tukey smoothing (repeated smoothing until convergence) (50) using R, and defined as the first time for which the distance is below 2 μm (only for trajectories starting at >3 μm, in order to be able to truly detect the polarization process). This threshold value was chosen looking at the distribution of plateau values for BSA- or *α*IgG-coated droplets. Tracking of the cell for analysis of centrosome orientation was performed by first obtaining a mask of the cell, from SirTubulin background cytoplasmic signal. This channel is used to create a mask of the cell on Fiji and find its center of mass. Briefly, the 3D stack is interpolated (to obtain an isotropic voxel), a background subtraction (based on a Gaussian filtered (radius=4) image of the field without cell, time=0) is applied. A Gaussian filter is applied on the resulting image (radius=2) to remove local noise and the cell is finally segmented using an automatic threshold (Huang). Advanced analysis of centrosome trajectories was performed by using the 3D cell contour generated on Fiji, and then computing the distance of the centrosome from the center of the cell, and the angle formed with the cell-droplet axis on Matlab, to merge this data with advanced nucleus analysis data. For experiments using Nocodazole, the centrosome was visualized by expressing eGFP-cent1, and tracked in the same way.

**Analysis of the Golgi Apparatus** was performed on Icy Bioimage analysis software (51). 3D masks of the Golgi apparatus and the droplet were obtained, and the average distance of the Golgi apparatus to the surface of the droplets was computed using a 3D distance map from the droplet. Characteristic times were extracted on single kinetic curves smoothed using 3R Tukey smoothing (repeated smoothing until convergence) (50) using R, and defined as the first time for which the distance is below 4 μm (only for trajectories starting at >5 μm, in order to be able to truly detect the polarization process). This threshold value was chosen looking at the distribution of plateau values for BSA- or *α*IgG-coated droplets.

**Analysis of the lysosomes** was performed on Icy Bioimage analysis software (51). 3D masks of the lysosomes and the droplet were obtained, and the average distance of all the lysosomes to the surface of the droplet was computed using a 3D Distance map from the droplet. Characteristic times were extracted on single kinetic curves smoothed using 3R Tukey smoothing (repeated smoothing until convergence) (50) using R, and defined as the first time for which the distance is below 3 μm (only for trajectories starting at >4 μm, in order to be able to truly detect the polarization process). This threshold value was chosen looking at the distribution of plateau values for BSA- or *α*IgG-coated droplets.

**Analysis of the Nucleus and detection of nuclear indentation** was performed using customs Fiji macros and Matlab software (available on request). B cell nucleus is bean-shaped and exhibits a marked invagination. To automatically detect the invagination at each time point, we interpolated the confocal images of the nucleus to obtain an isotropic voxel, segmented the nucleus and found the interpolating surface (isosurface function in Matlab). We smoothed the surface to reduce voxelization and computed the mean curvature at each vertex with standard differential geometry methods. We defined the invagination as the point with the minimal mean curvature obtained on this surface. *Ad hoc* correction based on nearest neighbor tracking is applied when several local minima are found (in nuclear that exhibit several lobes), the selected minimum is the nearest one to the point found in the previous frame. The orientation of the nucleus with respect to the Cell*_Center_*-Droplet_*Center*_ axis is quantified as the angle N*_indentation_*-Cell*_Center_*-Droplet_*Center*_.

## ACKNOWLEDGEMENTS

The authors thank M.Bolger-Munro for critical reading, H.Moreau, J.Delon, M. Théry, J.Husson and C.Hivroz for fruitful discussions, and acknowledge the Nikon Imaging Center@CNRS-Institut Curie, the PICT-IBiSA, Institut Curie, Paris, member of the France-BioImaging national research infrastructure, and the Cell and Tissue Imaging Platform - PICT-IBiSA (member of France–Bioimaging ANR-10-INBS-04) of the Genetics and Developmental Biology Department (UMR3215/U934) of Institut Curie (supported by the European Research Council (ERC EPIGENETIX N°250367)) for support in image acquisition, as well as the Recombinant Antibody Platform (TAb-IP) of Institut Curie. This work benefited from the technical contribution of the Institut Pierre-Gilles de Gennes joint service unit CNRS UAR 3750. The authors would like to thank the engineers of this unit for their advice during the development of the microfluidic setup.

PP and AMLD were supported by CNRS and INSERM, respectively. JP and LP were funded by the Ecole Doctorale FIRE-Programme Bettencourt, Université de Paris and the IPV scholarship program (Sorbonne Université), respectively. JF acknowledges funding from the Agence Nationale de la Recherche (ANR Jeune Chercheur PHAGODROP, ANR-15-CE18-0014-01).

## AUTHOR CONTRIBUTIONS

JP, AMLD and PP designed the study. JP set up and performed all the experiments, analyzed experimental results and prepared the figures. PP set up and performed analysis of centrosome and nucleus orientation. JP, LP and JF performed droplet formulation and characterization. OM designed the microfluidics chip and carried out preliminary experiments. JP, AMLD and PP wrote the manuscript. All authors approved the final version of the article.

## COMPETING INTERESTS

The authors declare that no competing interests exist.

## Supplementary Figures

**Fig. S1.**
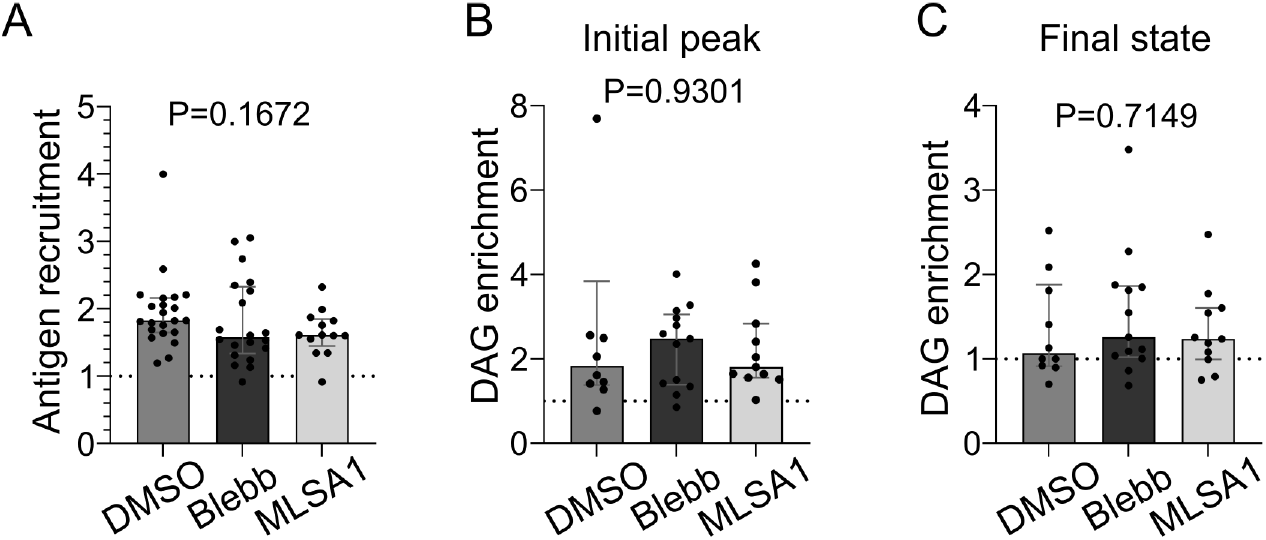
Myosin II merely regulates antigen recruitment and DAG signaling. (A) Plateau of antigen recruitment (average 25-30 min) (Median±IQR, DMSO N=22, p-nBlebb 20 μM N=20, MLSA1 1 μM N=13, 2 independent experiments, Kruskal-Wallis test). (B) Maximum (in 0-20 min) and (C) average final (25-30 min) DAG reporter enrichment (Median±IQR, DMSO N=10, p-nBlebb 20 μM N=13, MLSA1 1 μM N=11, 2 independent experiments, Kruskal-Wallis test).

## Notes

### Competing Interest Statement

The authors have declared no competing interest.

